# Anti-oxidant, anti-apoptotic, anti-hypoxic and anti-inflammatory conditions induced by PTY-2 against STZ-induced stress in islets

**DOI:** 10.1101/670364

**Authors:** Shivani Srivastava, Harsh Pandey, Surya Kumar Singh, Yamini Bhusan Tripathi

## Abstract

**Background and Aim:** Earlier assessment of *Pueraria tuberosa* tubers has shown anti-diabetic effects through incretin mimetic action and DPP-IV inhibition. The aim of this work was to further explore the protective role of aqueous extract of *Pueraria tuberosa* against streptozotocin (STZ)-induced pancreatic stress in rats.

**Methods:** Diabetes was induced with STZ (65 mg/kg body weight) in Charles foster male rats. After 60 days of STZ administration, animals with blood glucose levels > 200 g/dL were considered as diabetic. All the rats were later divided into three groups: Group-1 (STZ untreated normal rats), Group-2 (Diabetic control), and Group-3 (PTY-2 [50 mg/100 g bw treatment for next 10 days to diabetic rats). The rats were then sacrificed at the 10^th^ day of treatment accordingly.

**Results:** STZ treatment led to an increase in expression of MMP-9, Tnf α, HIF-1α, VEGF, IL-6, PKC ε, NF-kB, and Caspase-3. Reverse transcriptase Polymerase Chain Reaction (PCR), IHC and western blot analysis showed an increase in the expressions of superoxide dismutase (SOD) and Nephrin, and a decrease in the expressions of NF-kB, PKC ε, TNF-α MMP-9, HIF-1α, VEGF, Caspase 3 and IL-6 after 10 days of PTY-2 treatment.

**Conclusion:** The results show that PTY-2 favorably changed the expression of NF-kB, PKC ε, TNF α, MMP 9, HIF-1α, VEGF, IL-6, Caspase3, Nephrin and SOD in cases of STZ-induced pancreatic stress. Further evaluation of PTY-2 might be helpful in establishing its role in the management of diabetes mellitus.

**Highlights:**

1. PTY 2 act as a protective herbal drug against STZ induced islet stress.
2. PTY 2 upregulates protective and downregulates harmful markers.
3. This study composed of four pathway through which PTY 2 acts on pancreas.

**GRAPHICAL ABSTRACT:** Mechanism of action of PTY 2 against STZ induced islet stress.

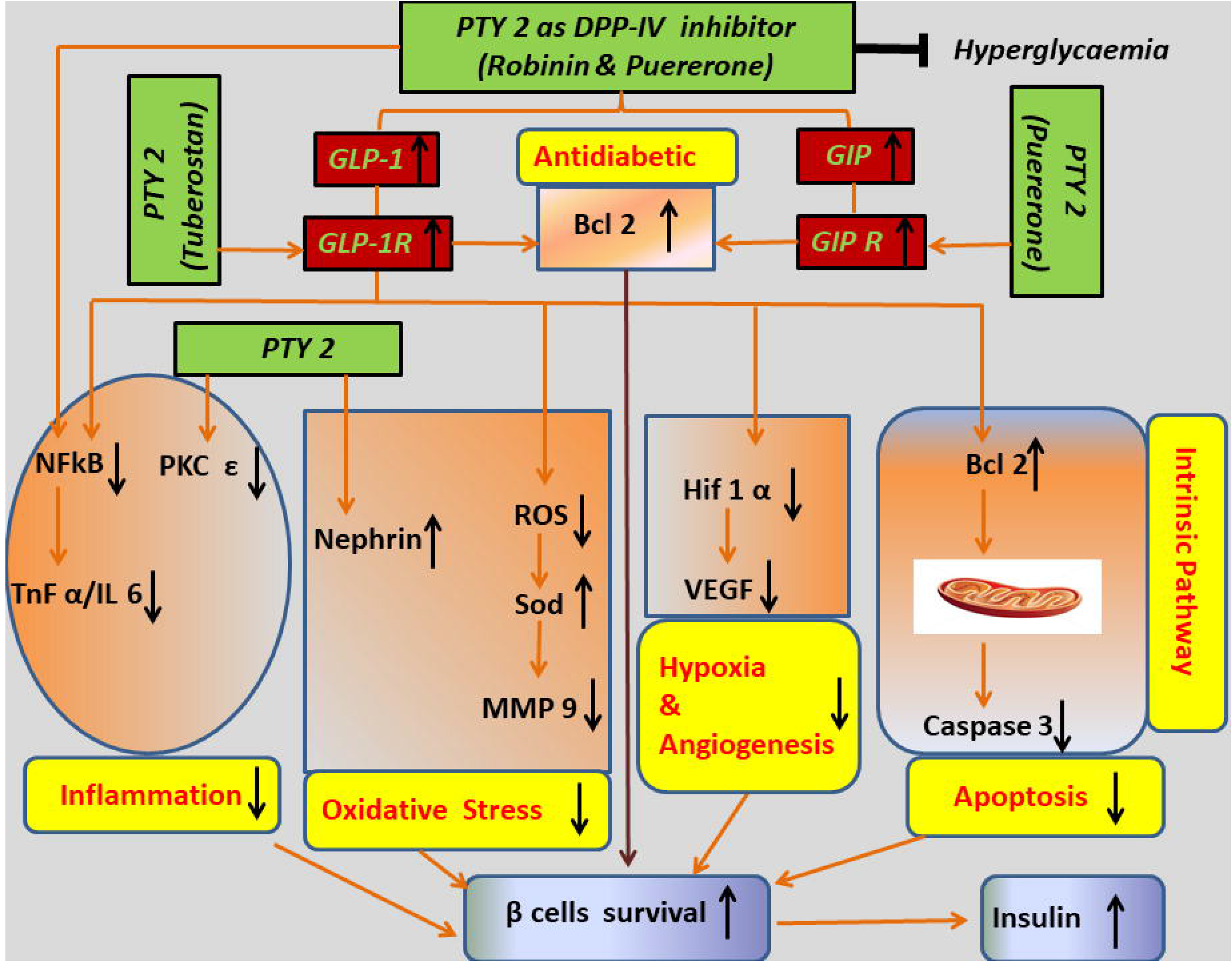

## Background

In addition, mortality and morbidity due to diabetes mellitus (DM) are also rising rapidly worldwide [1,2]. Type 2 DM (T2DM) increases the risk of acute pancreatitis by 1.5-3 folds, and the use of anti-diabetic drugs decreases this excess risk [2]. On the other hand, pancreatitis is one of the known risk factors for the onset of DM [3–5]. Additionally, the onset of DM can be a symptom of pancreatic cancer as the latter is more common among newly diagnosed cases of T2DM. Furthermore, long-standing DM can increase the risk of occurrence of pancreatic cancer [6–9]. Apart from the known etiological factors, pathological changes due to environmental factors or other unknown reasons can alter the gene expression and lead to diseases like DM, pancreatic cancer, acute or chronic pancreatitis [10,11]. Both *in-vivo* and *in-vitro* models are being developed to understand the mechanisms underlying the profile change in gene expression. Many synthetic drugs and herbal formulations have been developed for the prevention and treatment of DM.

The aqueous extract of one of the herbal products, *Pueraria tuberosa* tubers (PTY-2), is being evaluated for its protective role in STZ induced islet stress. In our earlier studies, we had evaluated the role of PTY-2 in animal models of streptozotocin (STZ)-induced DM and in normoglycemic rats [11–13]. PTY-2 has also been studied for its anti-inflammatory, antioxidant, nephroprotective, anti-hypertensive, anxiolytic and hypoglycemic properties [12,14–20]. The analysis of actions of PTY-2 shows that there must be interconnecting signaling pathways between anti-inflammatory, antioxidants and anti-diabetic genes for the effect of PTY-2. Taking our research forwards, we have attempted to study the signaling pathways to understand the role of antioxidant, antiapoptotic, anti-hypoxic and anti-inflammatory properties in providing a protective role in pancreatic damage.

Various markers like Nephrin, Superoxide dismutase (SOD), Hypoxia Inducible Factor-1α (HIF-1α Tumor Necrosis Factor-α (TNF-α), Matrix metalloproteinase 9 (MMP-9), Caspase-3, Nuclear factor kappa B (NF-kB), Vascular Endothelial Growth Factor (VEGF), Protein kinase C epsilon (PKC ε), and Interleukin-6 (IL-6) have been used to study the effect of drugs or herbal products on DM. Earlier studies have demonstrated increased levels of VEGF, TNF-α, MMP-9, IL-6, NF-kB, PKC ε, and HIF 1α in inflammatory conditions, vascular lesions and DM [21–25]. Excess generation of reactive oxygen species (ROS) and oxidative stress is one of the common etiological pathways in the development and progression of DM [26,27]. Nephrin, a member of immunoglobulin super family, is a surface receptor that is specifically expressed in kidney, brain and pancreas [28]. In the beta cells of islet of Langerhans of pancreas, nephrin plays an important role in beta cell survival signaling through the association with PI3-kinase in mouse islet β-cells and mouse pancreatic beta-cell line (βTC-6 cells) [29]. VEGF is a vital regulator of vascularization of islet cells, and the islet vascular system is critical for a normal section of insulin [30,31]. Genetic studies have shown that normal VEGF and vascularization are important for adult islet cell function and β cell mass [25]. The β cell-specific overexpression of VEGF causes rapid hypervascularization and hyperinnervation of the islet, leading to increased production of extracellular matrix components (ECM) [32]. Hence, we can say that increased amount of VEGF is responsible for defective angiogenesis. MMPs are a large family of endopeptidases, and these are produced by stromal and inflammatory cells. Pancreatic MMPs (especially MMP-9) induce inflammation, and serum MMP-9 levels are an assessment marker of severity of pancreatitis [33]. MMP-9 is usually involved in degradation and remodeling of ECM components and cellular migration [22,34,35]. NF-kB, a nuclear transcription factor, regulates the transcription of various genes involved in inflammation mediation [36]. The activation of NF-kB is an early pathological event in the development of insulin resistance [37]. TNF-α, an inflammatory marker, is rapidly produced intracellularly with the activation of NF-kB and is known to have effects on insulin transduction and obesity [21,38].The PKC ε belongs to the superfamily of isoforms of protein kinases. PKC ε is involved in the development of insulin resistance, and its inhibition is associated with the improvement in glucose homeostasis in animal models [39]. PKC ε has a strong presence in islet cells, acinar cells, and ductal epithelium [40]. Similarly, both, IL-6 and HIF-1α are also known to play a pro-inflammatory role in the mediation of acute pancreatitis and pancreatic cancer [23,24]. Hypoxia is an important cause of beta-cell loss and is measured by an increase in HIF-1α expression [41]. Various gene knockout experiments have shown that caspase-3 is involved in beta cell apoptosis and that *Casp^−/−^* are protected from the development of DM [42].

As *Pueraria tuberosa* has multiple medicinal properties with several beneficial compositions, we have studied the protective effect of its total water extract rather than on its individual components. Because *Pueraria tuberosa* contains many steroids, triterpenoid, glycosides, carbohydrates, alkaloids, flavanoid, tannin, protein and amino acids,e.g., daidzin, puerarin, puerarone, genistein, puetuberosanol, tuberostan, tuberosin, and puerarin 4’,6’-diacetate as the main constituents [12,14,43]. We planned to study the multitargeted protective effect of PTY-2 on the pancreatic damage among rats with STZ-induced stress.

## Research design and methods

### Materials

The antibody of rabbit IL-6 (23 Kda) (08310): SAB1408591, mouse monoclonal VEGF(21 Kda) (JH-121): sc-57496, NF-κB p65(D14E12) XP^®^ Rabbit mAb #8242, rabbit polyclonal PKC-ε (SAB1300094), mouse monoclonal β-actin (A2228), monoclonal Hif-1α (H6536-100 UG), rabbit monoclonal Caspase 3 (CASP 3 [D175] invitrogen), Rabbit monoclonal antibody to MMP 9 and monoclonal anti-rabbit IgG (◻-chain specific)-peroxidise (A1949), pre-stained protein ladder (from Hi-Media Pvt. Ltd, Kolkata, India) along with PVDF membranes (from Millipore, catalog no. IPVH20200) were used for western blotting. STZ-S0130 was bought from Sigma-Aldrich, St Louis, USA. For RT-PCR, Trizol (Himedia, Pvt. Ltd, Kolkata, India), cDNA Kit (Fermentas), and Taq-polymerase (Genaxy Scientific Pvt.Ltd) were used.

### Sample preparation

*Pueraria tuberosa* was purchased from Ayurvedic Pharmacy, Banaras Hindu University. Its authenticity has already been ascertained in our previous research [44]. We extracted 30 g powder with eight volumes of distilled water. When the volume was reduced to ¼th, it was filtered with cloth. The total yield of *Pueraria tuberosa* water extract (PTY-2) obtained by this process was 30%.

### Study Design

The protocol was approved by the Institute Ethical Committee (Dean/2015/CAEC/1266), Institute of Medical Sciences, Banaras Hindu University. After overnight fasting, Charles foster male rats of the same age group with body weight in the range of 120-130 grams were injected STZ (65 mg/kg body weight). STZ was prepared in chilled and fresh citrate buffer of pH 4.5. The blood glucose levels were checked using strips (Dr.Morepen) on the 5^th^ day. Rats with blood glucose levels > 200 mg/dL were placed under diabetic group. In order to induce severe diabetes, we left the rats (three rats per cage) for 55 days. On the 61^st^ day, we divided the rats into three groups (n=6): Group-1 (STZ untreated rats, i.e., age-matched normal control), Group-2 (diabetic control), and Group-3 (PTY-2 at 50 mg/100 g bw treatment for 10 days to diabetic rats). The rats were then sacrificed after 10 days of treatment. The pancreas was isolated and rinsed with PBS. Then, these were cut into two parts; one for histology (preserved in 10% formaldehyde) and the other was first crushed in liquid nitrogen and then stored in −80°C freezer for molecular study.

### Reverse transcription-polymerase chain reaction (RT-PCR)

RNA was extracted using Trizol reagent from about 50 mg of pancreatic tissue with a homogenizer. Then 5 μg of total RNA was reverse-transcribed with superscript II RNase H-reverse transcriptase (RT) using random hexamers according to the instructions provided by manufacturers (Fermentas Pvt. Ltd.). For SOD, 2 μl c-DNA, 0.2 mmol/L deoxynucleotide triphosphates (dNTPs), 1.5 mmol/L MgCl_2_, 0.5 μmol/L of each primer, 2.5 μl 10X PCR buffer and 1U Taq DNA polymerase were used. For Nephrin, 1 μl c-DNA, 0.2 mmol/L dNTPs, 1.5 mmol/L MgCl_2_, 1.2 μmol/L of each primer, 2.5 μl 10X PCR buffer and 1U Taq DNA polymerase were used. For matrix metallopeptidase 9 (MMP 9), 2 μl c-DNA, 200 umol/L dNTPs, 1.5 mmol/L MgCl_2_, 0.4 μmol/L of each primer, 2 μl 10X PCR buffer and 2.5 U Taq DNA polymerase were used. For Tnf α 1 μl c-DNA, 200 umol/L dNTPs, 1.2 mmol/L MgCl_2_, 0.6 μmol/L of each primer, 2 μl 10X PCR buffer and 2U Taq DNA polymerase were used. For glyceraldehyde 3-phosphate dehydrogenase (GAPDH), 0.1 μmol/L of each primer was used. The optical density of each expression was determined via alpha imager (Bio-Rad) and presented as the ratio against GAPDH. All RT-PCR experiments were performed in triplicates (Table 1).

**Table 1.**
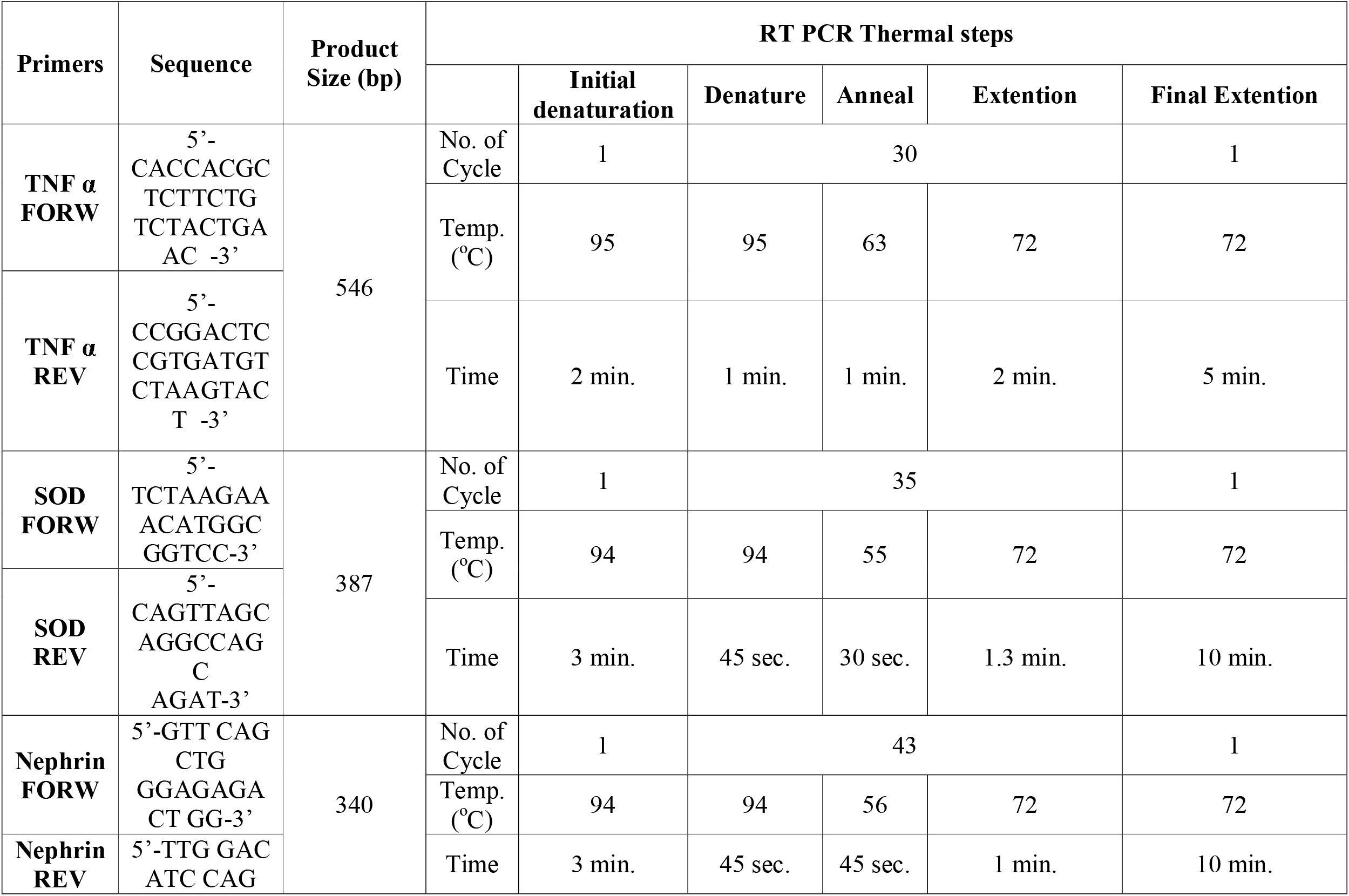
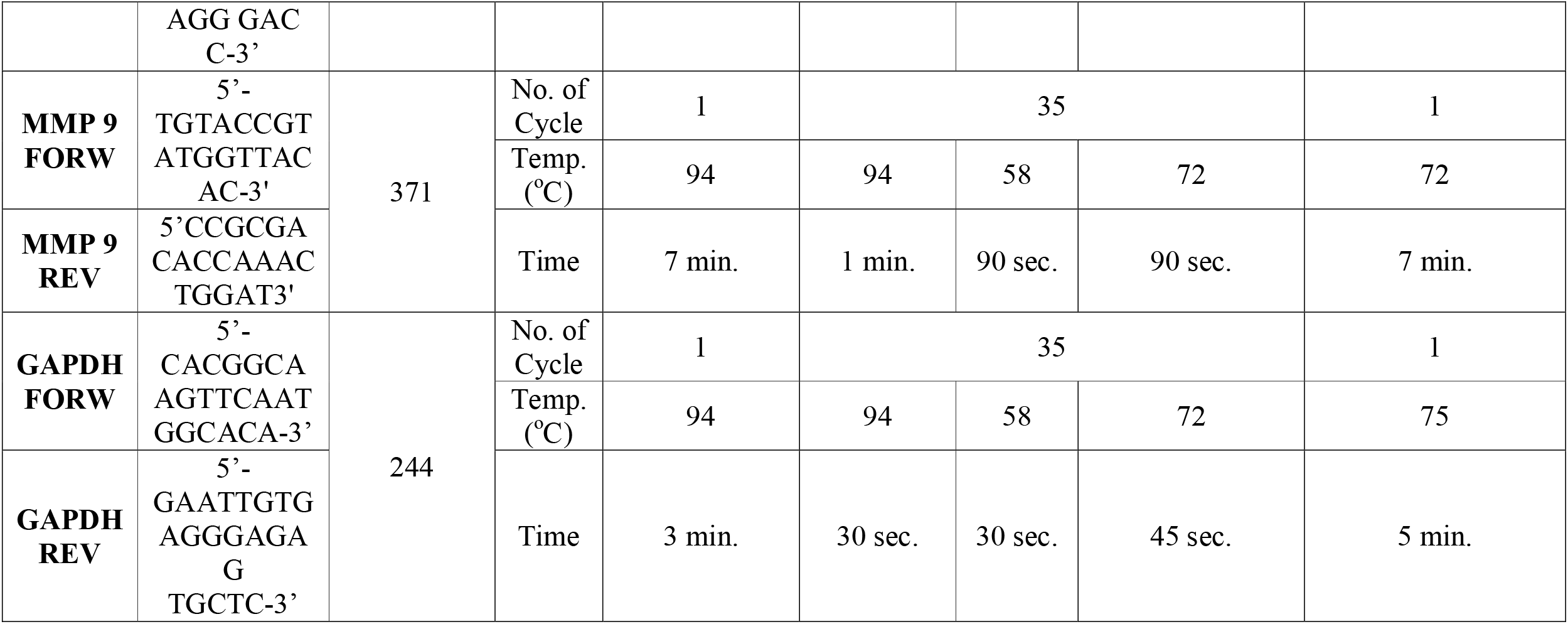
Details of PCR primer sequences, product size and thermal steps for expressions of TNF α, SOD, Nephrin, MMP 9 and GAPDH.

### Western blot analysis

Washed pancreatic tissue (10% ice-cold PBS) was homogenized with chilled lysis buffer (50 mMTris pH 7.6, 150 mM NaCl, 1 mM EDTA, 1 mM EGTA, 1% triton, 0.1% SDS, 1 mM sodium orthovanadate, protease inhibitor cocktail and 1 mM PMSF). The homogenate was then centrifuged at 12,000 rpm at 4°C for 30 min. The protein estimation was done by Bradford method according to which 40μg proteins with loading dye were separated in the polyacrylamide gel. The gel was then electro-transferred to PVDF membranes in transfer buffer (10X Tris-glycine-methanol and SDS-PAGE buffer) to stay overnight at 4°C at 45 V. The next day, PVDF membrane was blocked with 5% non-fat milk powder. The membrane was then incubated overnight with primary antibody diluted in TBST [IL-6 (1:1000), PKC ε (1:500), VEGF (1:1000), NF-kB (1:1000) and HIF 1α (1:1000) &housekeeping gene β-actin (1:500)]. Then, on the next day, the blots were incubated with secondary antibody (1:50000) in TBST for one hour. Protein expression was detected through enhanced chemiluminescence (ECL) in LAS 500 Image Quant system (Wipro GE Healthcare, Hong Kong). The quantification was done by Alpha imager (Protein Simple 3001 California, USA). The experiments were done in triplicates.

### Immunostaining

The paraffin sections of pancreas were treated with Xylene for 10 minutes to remove paraffin. The sections were rehydrated through 90%, 70% alcohol, and water by putting them for 5 minutes in each. Antigen retrieval was done by putting the citrate buffer dipped slides in EZ Retrieval System V.3 (Bio Genex). Sections were washed twice in citrate buffer and two times in 1X PBS (130 mM NaCl, 7 mM Na_2_HPO_4_, 3 mM KH_2_PO_4_, pH 7.4) for 10 minutes each, following which the sections were incubated in blocking solution [0.1% Triton X-100, 0.1% BSA, 10% FCS, 0.1% sodium deoxycholate and 0.02% Thiomersal (an anti-fungal agent), in 1X Phosphate Buffered Saline (PBS)] for 2 hours at room temperature and then transferred in rabbit primary antibodies, for overnight at 4oC. Tissues were washed in PBST (0.1% triton X in 1XPBS) with three changes of 10 minutes each. After the washing, the sections were incubated with anti-mouse-AF546 and anti-rabbit-AF488 (Invitrogen, USA) secondary antibody for 2 hours at room temperature. Sections were washed in PBS with Tween 20 (PBST) with three changes for 10 minutes each, counterstained with DAPI (1 μg/ml DAPI in 1XPBS), mounted in DABCO and examined under Zeiss LSM510 Meta confocal microscope. Image analysis was done by using Zen Black (2012) software.

### Statistical analysis

One-way ANOVA test followed by post hoc analysis with Dunnett’s test was done for each experiment. All results were expressed as means ± SD. Statistical significance was taken at *p* ≤ 0.05.

## Results

### PTY2 response to pancreatic inflammation

#### mRNA EXPRESSIONS

As compared to normal rats, the STZ-treated diabetic group showed a significant increase in TNF α in pancreatic tissue, whereas the PTY-2 treatment significantly decreased the TNF α expression as compared to diabetic control and increase as compared to normal. The MMP-9 expression also increased significantly in diabetic control as compared to normal rats, and there was a significant decrease after 10 days of PTY 2 treatment. On the contrary, both SOD and Nephrin expression decreased significantly in diabetic control rats as compared to normal. However, the PTY 2 treated group showed a significant increase in SOD expression as compared to diabetic control and a significant decrease as compared to normal. The Nephrin expression in PTY 2 treated rats increased significantly as compared to both normal and diabetic control (Fig. 1 (a) and (b))

**Fig. 1.**
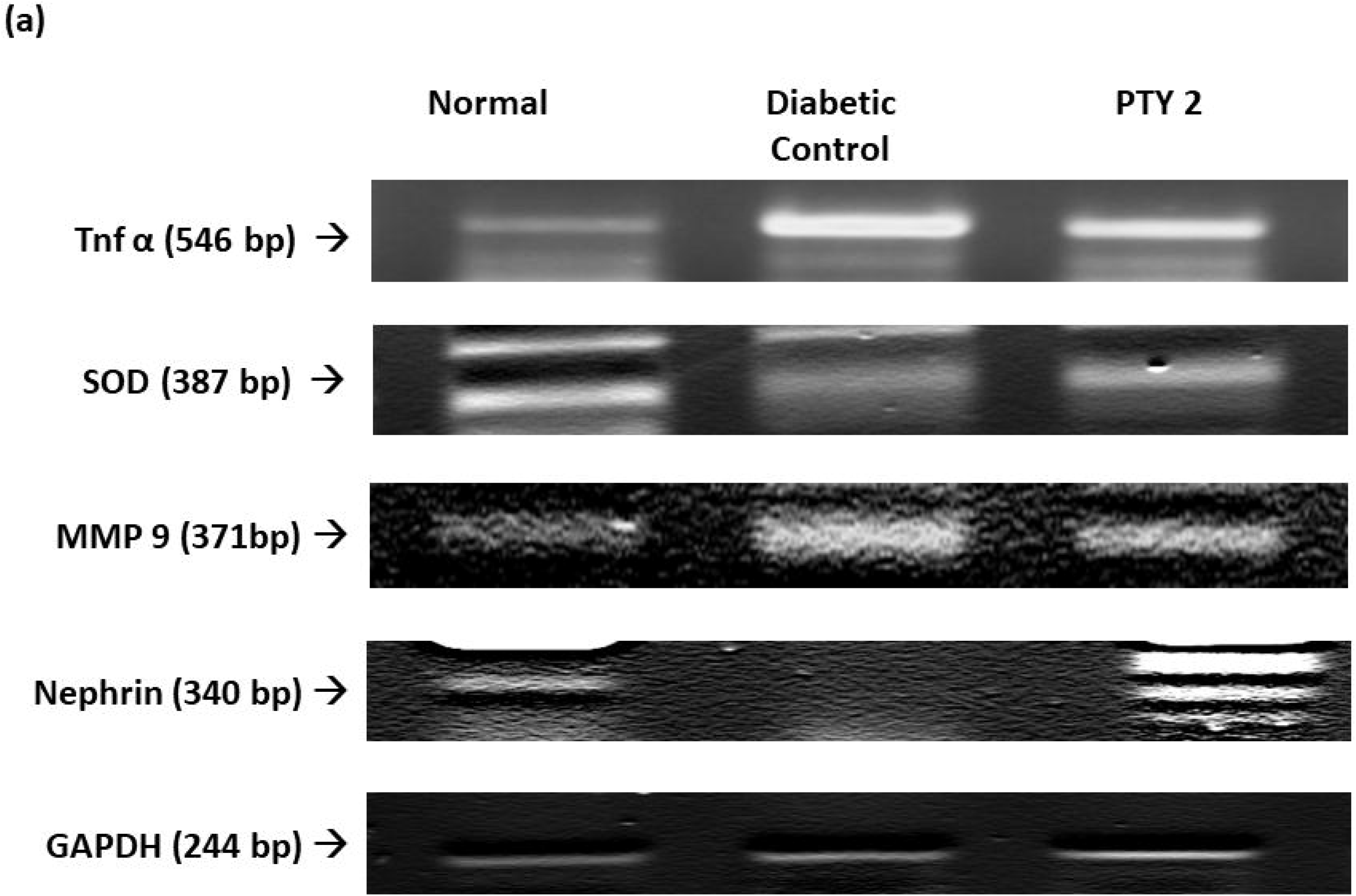
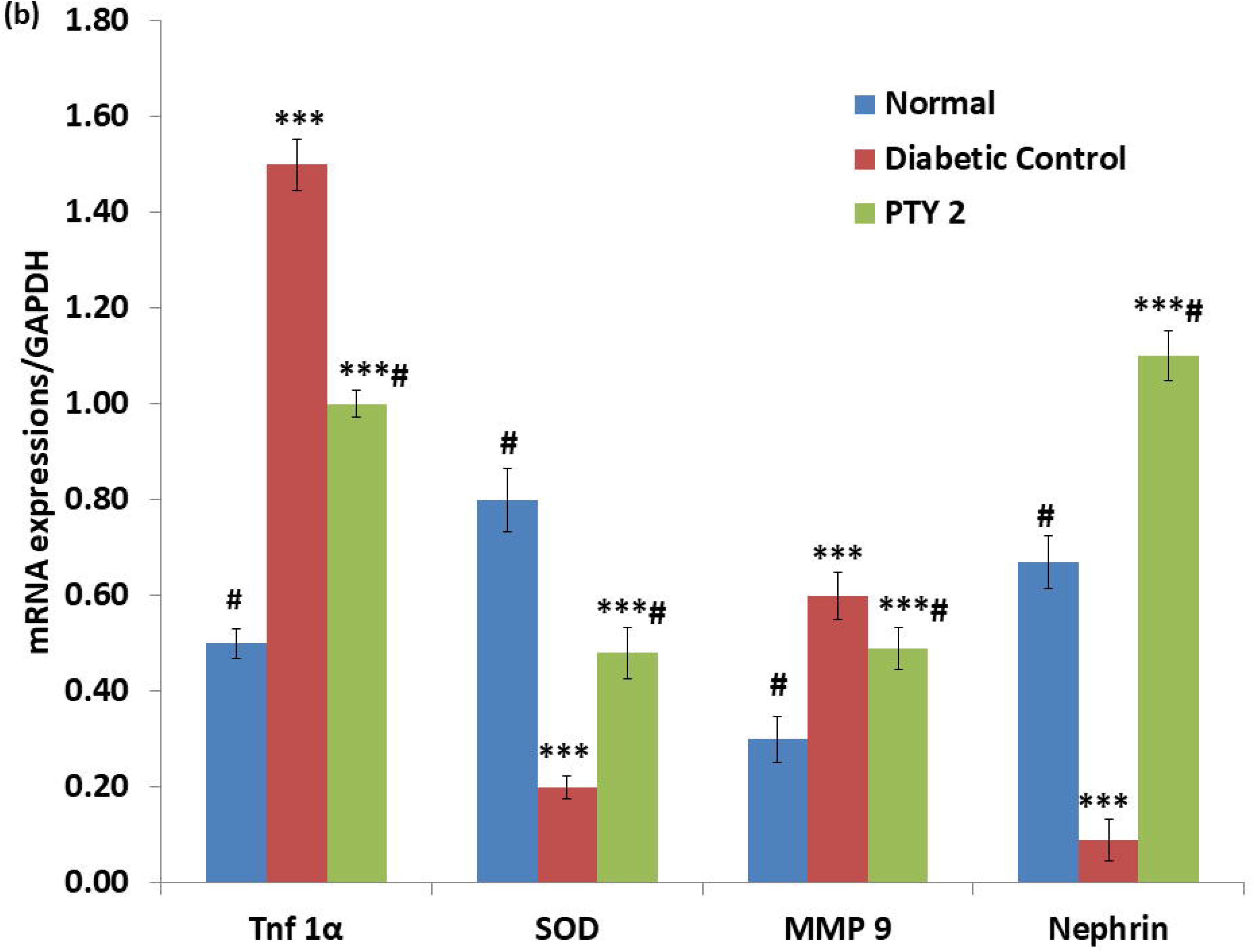
**a)** mRNA expressions to investigate the effect of PTY-2 on STZ induced pancreatitis; **(b)** Desitometry analysis of RT-PCR product. Each value represents mean ± SD (n=6); \*\*\**P*< 0.05 compared with group 1, # *P* < 0.05 as compared with group 2.

These results clearly indicate that in chronic diabetes, there is significant increase in free radicals/ stress accompanied by increase in pancreatic inflammation. Treatment with PTY 2 significantly reversed all these changes. Thus, any severe complications of severe diabetes like pancreatitis could be prevented by using PTY 2 as medicinal supplement.

#### PROTEIN EXPRESSIONS

##### Western blot

For further validation, the protein expressions responsible for the induction of oxidative stress, hypoxia, apoptosis and inflammation of pancreatic tissues were estimated. The expressions of NF-kB, PKC ε, HIF-1α, VEGF and IL-6 were significantly increased in diabetic control as compared to normal rats. However, PTY 2 treatment significantly decreased all these expressions (Fig. 2 (a) and (b)).

**Fig. 2.**
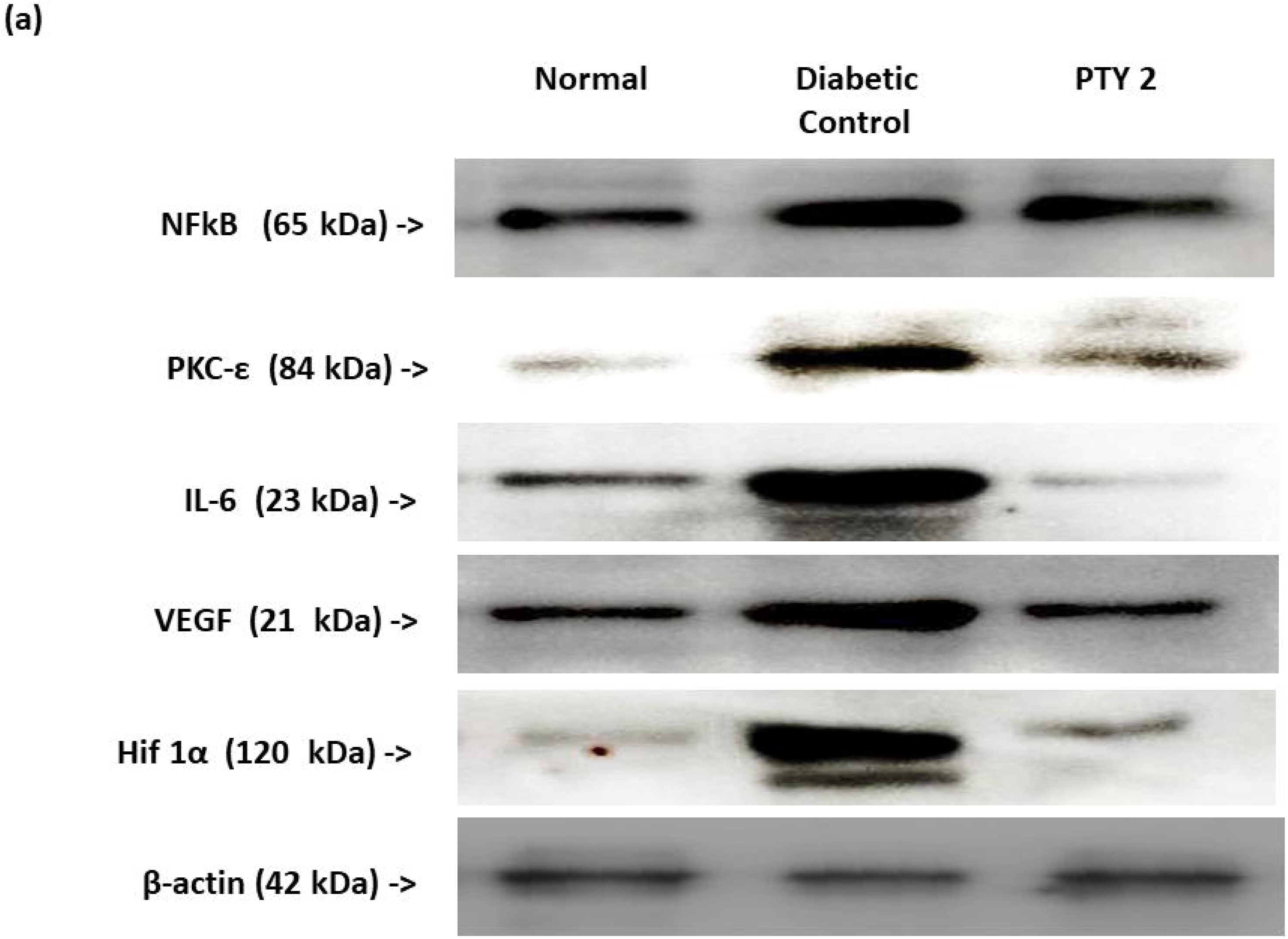
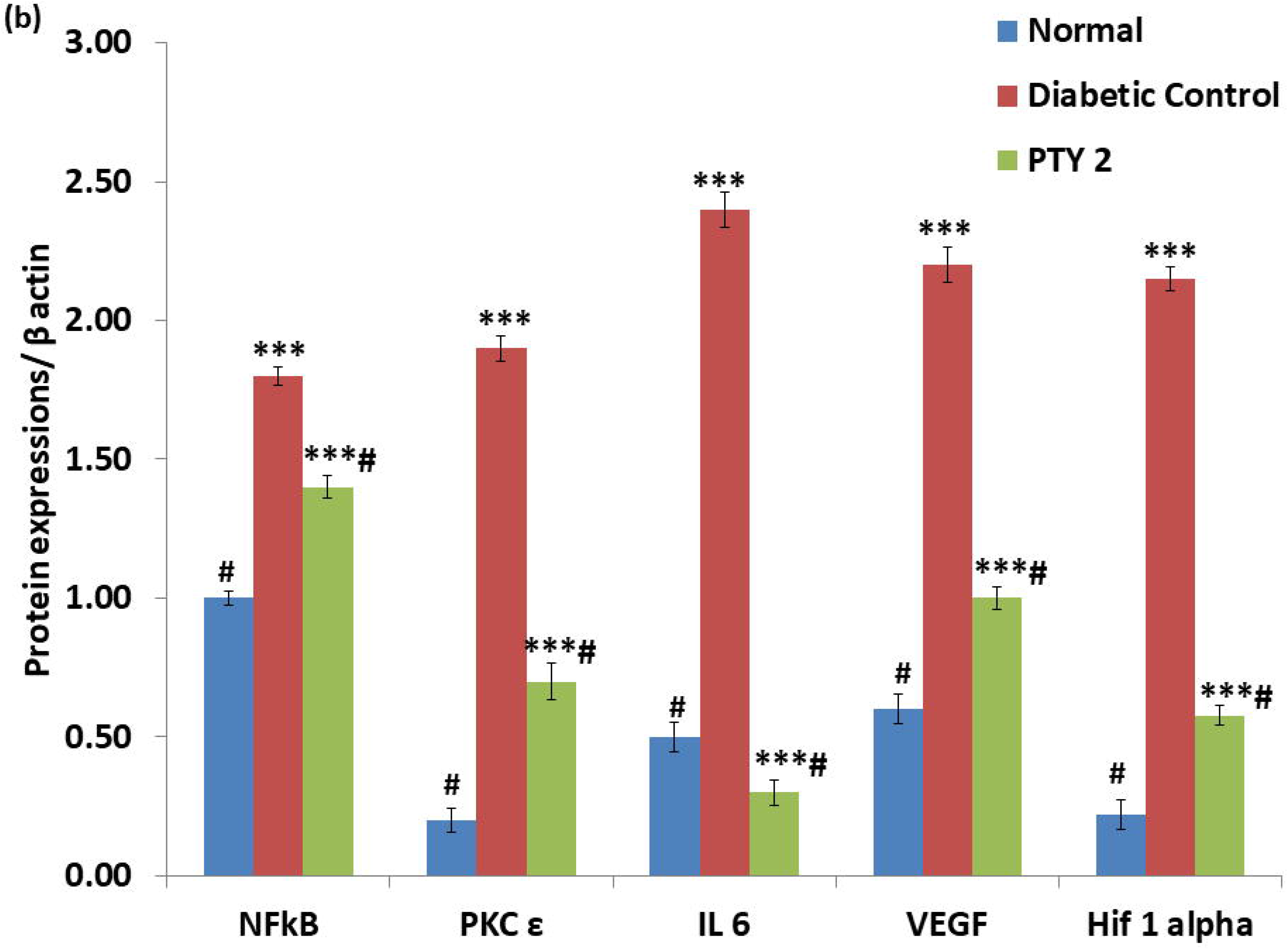
**a)** Protein expressions to investigate the effect of PTY-2 on STZ induced pancreatitis; **(b)** Desitometry analysis of western blot product. Each value represent the mean ± SD (n=6); *** *P* < 0.05 compared with group 1, # *P* < 0.05 compared with group 2.

##### Immunohistochemistry

The expressions of MMP-9, HIF-1α, VEGF, IL-6, PKC ε, NF-kB and Caspase-3 were significantly enhanced in diabetic control islets. The hazardous effects of STZ were down-regulated by 10 days of PTY-2 treatment. The Caspase-3, PKC-ε, HIF-1α and NF-kB expressions decreased significantly in PTY 2 treated group as compared to diabetic control and increased significantly as compared to the normal. The expressions of MMP-9, IL-6 and VEGF in PTY 2 treated group decreased significantly as compared to diabetic control, but non-significant to normal rats (Figures 3, 4, 5 & 6).

**Fig. 3.**
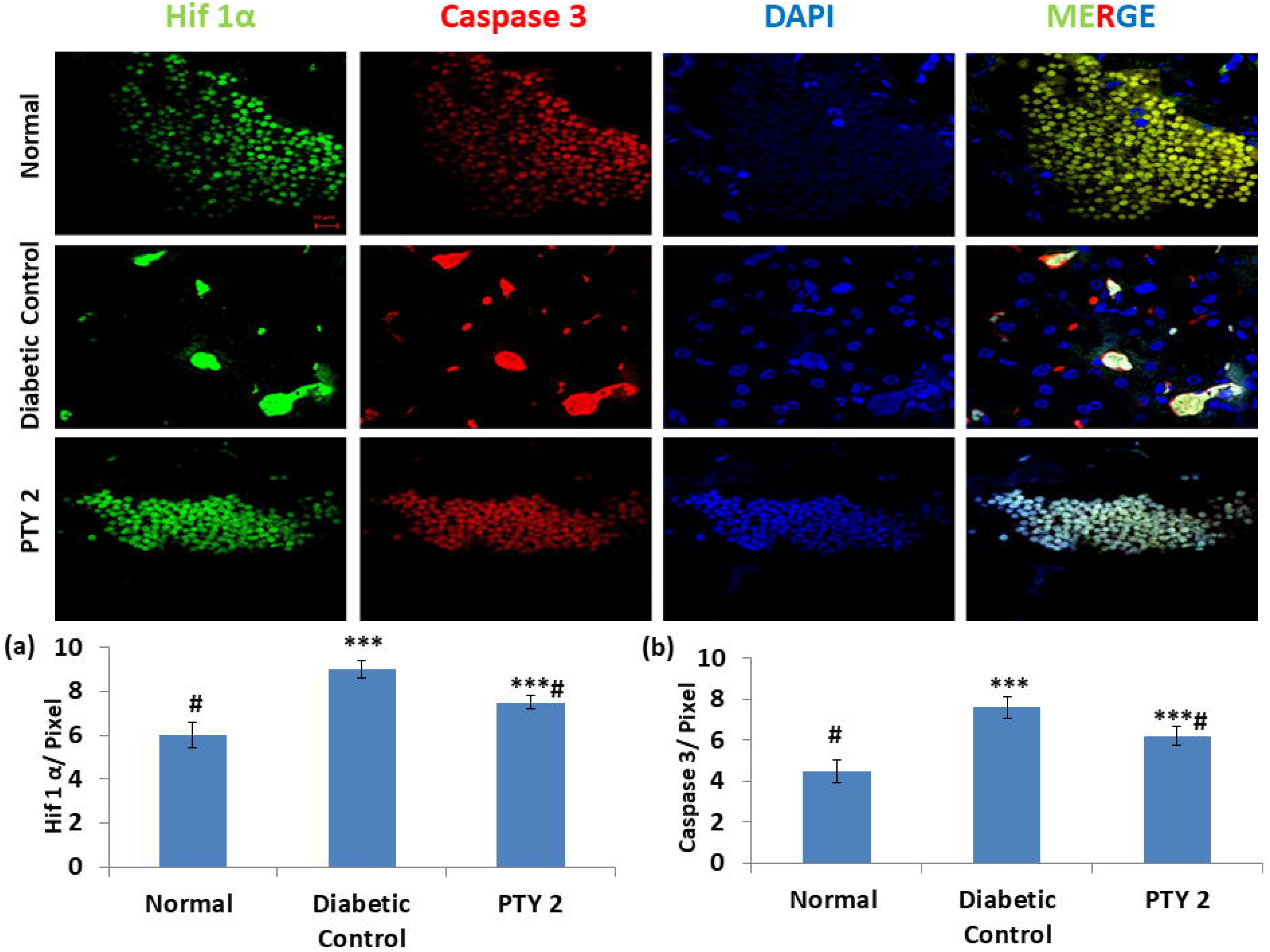
Immunohistochemistry analysis showed the effect of PTY-2 on the expression of **(a)** HIF-1α (green) and **(b)** Caspase 3 (red) in the islets of normal, diabetic control, and PTY-2-treated rats’ pancreatic tissues. Both the expressions were merged with DAPI (blue). In comparison to diabetic control, PTY-2 down regulated the expression of both HIF-1α and Caspase 3. The image was taken at 63X magnification. Scale bar was 20 μm. The intensity was measured in pixel values. Each value represent the mean ± SD (n=6); *** *P* < 0.05, compared with group 1, # *P* < 0.05, compared with group 2.

**Fig. 4.**
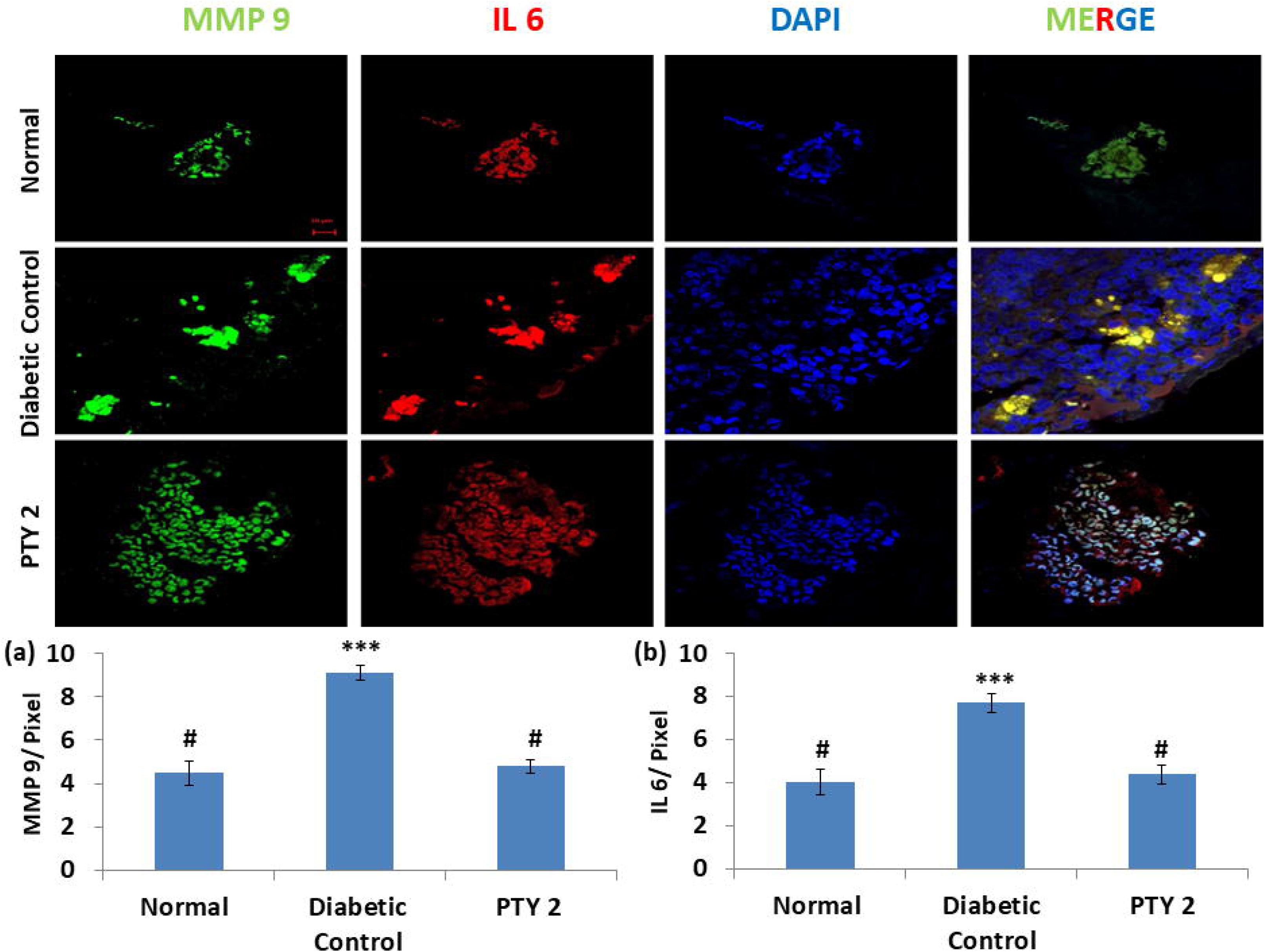
Immunohistochemistry analysis showed the effect of PTY-2 on the expression of **(a)** MMP-9 (green) and **(b)** IL-6 (red) in the islets of normal, diabetic control, and PTY-2-treated rats’ pancreatic tissues. Both the expressions were merged with DAPI (blue). In comparison to diabetic control, PTY-2 down regulated the expression of both MMP-9 and IL-6. The image was taken at 63X magnification and scale bar was 20 μm. The intensity was measured in pixel values. Each value represent the mean ± SD (n=6); *** *P* < 0.05, compared with group 1, # *P* < 0.05, compared with group 2.

**Fig. 5.**
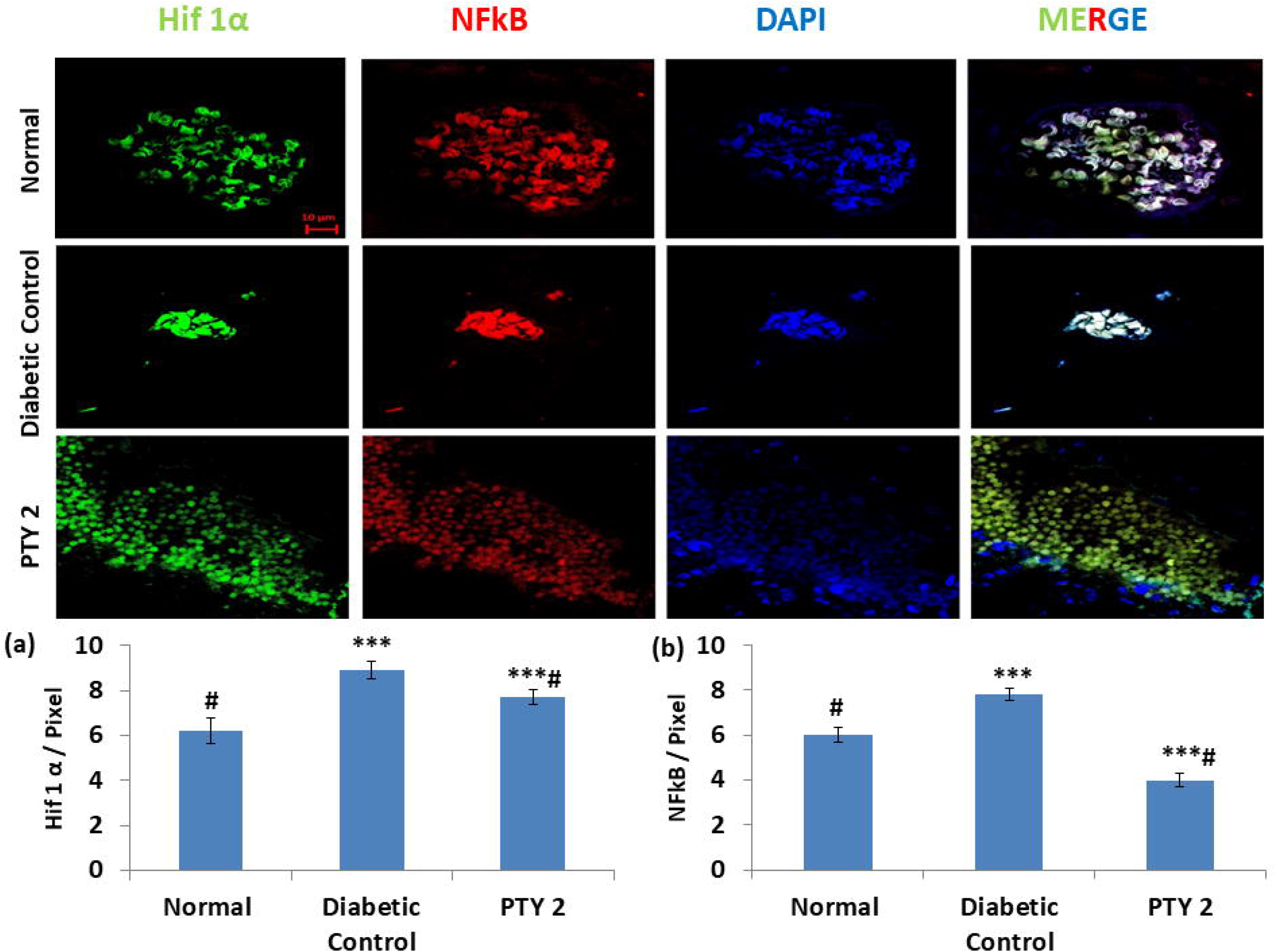
Immunohistochemistry analysis showed the effect of PTY-2 on the expression of **(a)** HIF-1α (green) and **(b)** NF-kB (red) in the islets of normal, diabetic control, and PTY-2-treated rats’ pancreatic tissues. Both the expressions were merged with DAPI (blue). In comparison to diabetic control, PTY-2 down regulated the expression of both HIF-1α and NF-kB. The image was taken at 63X magnification and scale bar was 20 μm. The intensity was measured in pixel values. Each value represent the mean ± SD (n=6); *** *P* < 0.05, compared with group 1, # *P* < 0.05, compared with group 2.

**Fig. 6.**
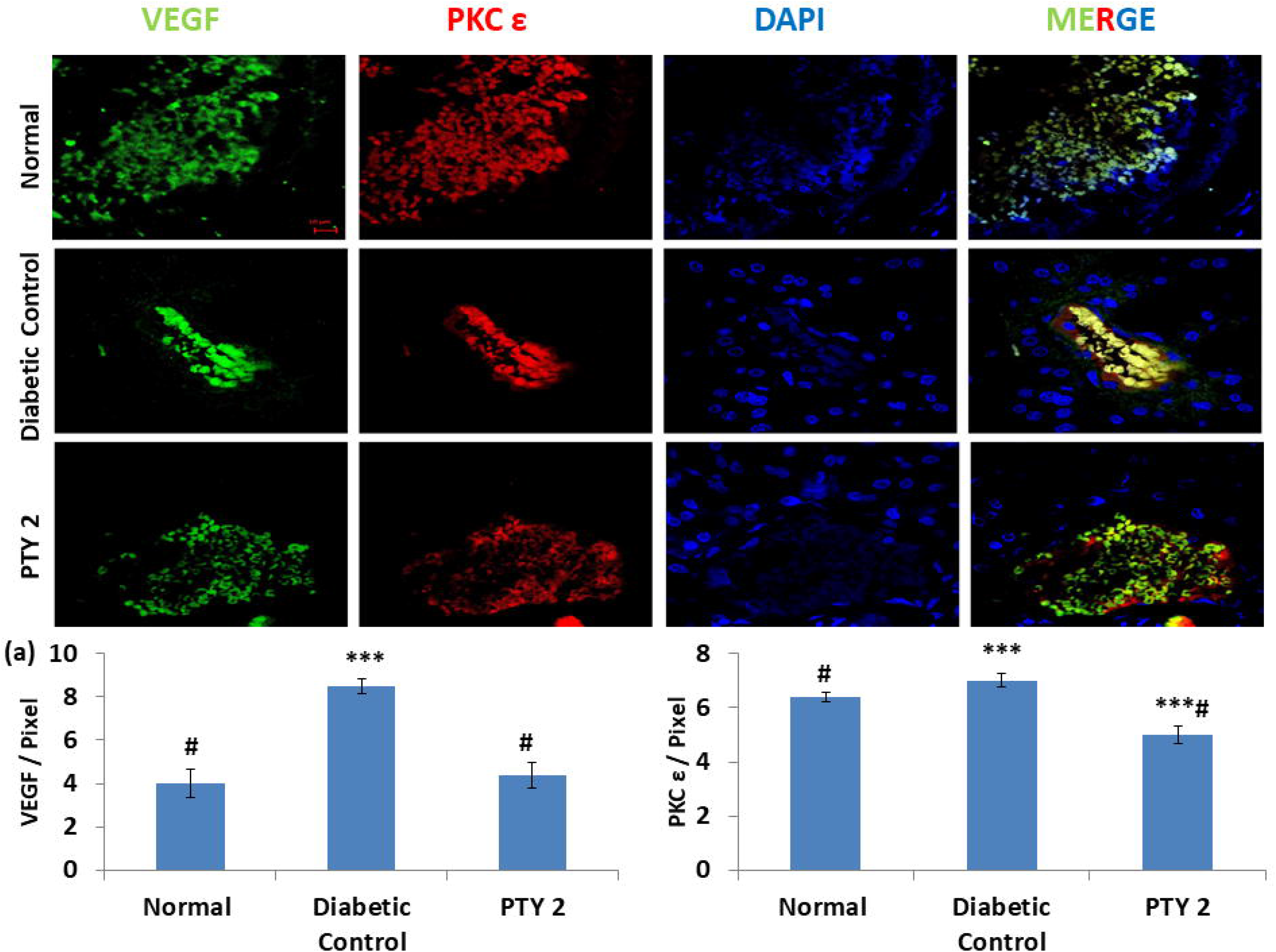
Immunohistochemistry analysis showed the effect of PTY-2 on the expression of **(a)** VEGF (green) and **(b)** PKC ε (red) in the islets of normal, diabetic control, and PTY-2-treated rats’ pancreatic tissues. Both the expressions were merged with DAPI (blue). In comparison to diabetic control, PTY-2 down regulated the expression of both VEGF and PKC ε. The image was taken at 63X magnification and scale bar was 20 μm. The intensity was measured in pixel values. Each value represent the mean ± SD (n=6); *** *P* < 0.05, compared with group 1, # *P* < 0.05, compared with group 2.

## Discussion

The results of our study showed that, compared to the diabetic control group, the PTY-2 group had a favorable change in the expressions of biomarkers as assessed by RT-PCR, Western blot and IHC techniques. Analysis showed that STZ increased the expressions of MMP-9, TNF- α, HIF-1α, VEGF, IL-6, PKC ε, NF-kB and Caspase-3 in diabetic control. But these expressions were significantly decreased by the treatment with PTY-2 for 10 days. Expression of SOD and Nephrin were significantly decreased among diabetic control rats, and these increased significantly after the administration of PTY-2. Both RT-PCR and western blot estimation showed a favorable change in expressions of the biomarkers for oxidative stress and inflammation.

Our earlier research had focused on the antidiabetic role of PTY-2 in STZ-induced diabetic model. In those studies, we found that PTY-2 has an effect on inflammation (Pandey et al., 2013), hyperglycemia and incretin pathways. Initially, the results showed that PTY-2 had hypoglycemic action because of inhibition of dipeptidyl peptidase-IV (DPP-IV) activity. In further research, we evaluated the effect on incretin receptor (GLP-1R and GIP-R), and insulin. The results showed that PTY-2 treatment led to a significantly higher increase in GLP-1 and GIP levels and a significant decrease in glucose concentrations after PTY-2 treatment (50 mg/100 g body weight) for 10 days. In the second study of chronic DM induced with STZ among rats,, there was also a significant decrease in DPP-IV activity and an enhanced basal insulin concentration in PTY-2 treated diabetic rats. Additionally, there was an increase in the number of islet cells and a significant increase in protein expression of insulin and Bcl-2. According to *in silico* study in our lab, Puerarone and Robinin were the two most effective phytochemicals for DPP-IV inhibition, and tuberostan and puererone identified as the active component against GLP-1 and GIP receptors. Moreover, an *in-vivo* experiment showed that anti-inflammatory property of *Pueraria tuberosa* might be because of the scavenging of free radicals by increase in activity of SOD, and decrease in C-reactive protein levels [15].

Moving in the same direction of evaluation of the role of PTY-2 in DM, we studied the effect of PTY-2 on the markers of oxidative stress, hypoxia, apoptosis and inflammation, which are known to play a significant role in the development and progression of pancreatitis and DM. We evaluated the change in expressions of mediators of inflammation and oxidative stress after STZ treatment followed by administration of PTY-2 for 10 days. The immunohistochemical, PCR and western blot analysis of our study showed an increase in the expression of MMP-9, TNF αHIF-1α,VEGF, IL-6, PKC ε, NF-kB and Caspase-3 as well as reduced expression of Nephrin and SOD after 60 days of STZ treatment in group 2 (Figures 1,2, 3, 4, 5, 6).

To understand and assimilate the results, the pathological changes in DM along with the proposed mechanism of action of PTY-2 according to our study have been diagrammatically shown in Graphical abstract. DM is a manifestation of abnormal metabolism and transport of glucose, and may be associated with a decrease in insulin secretion and presence of insulin resistance. Further on, this leads to hyperglycemia and an increase in the release of free fatty acids (FFAs)[45]. All these inter-related steps decrease β-cell function and number, increase oxidative stress, induce apoptosis and endoplasmic reticulum stress, and increase the release of inflammatory cytokines [45]. FFAs are known to induce the release of various interleukins, including IL-6, which further increase the release of free radicals and activate caspases. As shown in graphical abstract, NF-κ B, PKCε, TNFα, and IL-6 are mediators of inflammation, and the administration of PTY-2 in our study was associated with a decrease in the expression of all these mediators. With the progression of diabetes, the balance between the pro-inflammatory and anti-inflammatory or protective mediators is disturbed [45]. The increase in oxidative stress, measured by a decrease in SOD activity and an increase in ROS, was also decreased by PTY-2 administration. Other mediators of oxidative stress, i.e., MMP-9 and VEGF, were also decreased. Hypoxia, which is measured with HIF 1α, is also an inducible factor of DM and was decreased by PTY-2 administration. Apoptosis, one of the critical pathological changes, is mediated by an increase in activity of caspases [42]. PTY-2 led to a decrease in caspase-3 expression. Taking into consideration of our earlier evidence, it can be proposed that PTY-2 acts as a DPP-IV inhibitor, potentiates GLP-1 and GIP [13] mediated responses, and decreases inflammation, oxidative stress, apoptosis and hypoxia. GLP-1 agonists showed an inhibition of pro-inflammatory mediators in DM and other inflammatory conditions as well, in addition to their glucose-lowering potential [46,47].

MMP-9, one of the markers estimated in our study, deteriorates the inflammatory condition as it causes vascular injury, increases migration and cellular invasion by inflammatory cells [22,33]. Both animal and human studies have shown an increase in MMP-9 expression in pancreatitis [48,49]. MMP-9 acts as a diabetogenic factor by increasing proteolytic cleavage of insulin and generation of immunodominant insulin peptides [48,49]. Similar to our results, earlier studies have also shown an induction/increase of MMP-9 activity in STZ-induced models of DM [22]. It is assumed that hyperglycemia induced oxidative stress induces the expression of MMP-9 in pancreas, and this can be counterbalanced with GLP-1 agonists [22,50]. However, MMP-9 along with other paracrine factors is required for normal islet matrix turnover in the pancreatic islets [51]. IL-6, another inflammatory mediator, is also known to perform both inflammatory and protective roles [45]. In type 1 DM, IL-6 maintains the balance of cytokines; however, it is related with both initiation and progression of insulin resistance in T2DM [45]. However, PKC ε inhibition/deletion is associated with an improvement in glucose homeostasis [39]. In a previous study, when the Psammomys (sand rats) were fed with high energy diet, they developed insulin resistance mimicking T2DM. Treatment with PKC ε abrogated peptides and prevented insulin resistance, hyperinsulinemia and pancreatic beta cell loss. It shows that the enhanced expression of PKC ε in T2DM is associated with beta cell loss [52]. In another study with culture of lipid-treated islets isolated from PKCε knockout (PKCεKO) mice, there occurred amplification of GSIS (glucose-stimulated-insulin-secretion), reinforcing the benefit of inhibition of PKCε [39]. Our results also showed a significant increase in the diabetic group, followed by a significant decrease in the PTY-2 group.

Pro-inflammatory and pro-apoptotic cytokines like IL-1β are involved in the development and progression of insulin resistance via the decrease in the number and functioning of β cells. NF-κB is a transcription factor for mediating the cellular responses of inflammatory cytokines like IL-1β. The activation of NF-κB leads to a permanent decrease in the expression of proteins of β cells [45]. NF-kB pathways control cellular proliferation, inflammation, and immune responses through signal transduction [21,53]. The activity of NF-kB is increased in acute pancreatitis, and the longer duration of increased activity is associated with chronic disease [54–57]. Sitagliptin, a DPP-IV inhibitor used among T2DM patients, has shown anti-inflammatory action through the inhibition of NF-κB, inflammatory cytokines and cell apoptosis [58]. Mice deficient in NF-κB have shown to be resistant to STZ-induced diabetes [59].

In addition to the inter-relationship among MMP-9, IL-6, ROS, and NF-kB pathway, TNF-α is also a part of this pathological process [60]. The effects of TNF-α are mediated through the activation of NF-kB pathway. A decrease in IL-1action and an increase in the expression of TNF-α gene can increase the risk of onset and progression of the risk of DM [21]. Although the administration of TNF-α to animals is associated with insulin resistance and the regulation of TNF-α levels can improve insulin sensitivity, the status of TNF-α as a drug target for DM is still being evaluated. This might be possible with more understanding of the inter-relationships of the mediators in the pathogenesis of DM [61].

Hyperglycemia also leads to the destabilization of HIF-1α, which is responsible for the regulation of the cellular responses to hypoxia. [62,63]. Hypoxia is an important cause of apoptosis and beta cell loss, and HIF-1α is an important indicator of beta cell loss [64]. It is known that GLP-1R agonists (Extendin-4) improve islets survival through the activation of transcription factor, cAMP response element binding protein (CREB). A combination of CREB and Extendin-4 exerted enhanced anti-apoptotic action in cultured islets against hypoxia and cytokines. In an early phase, HIF-1α comes as a metabolic adaptation, but its prolonged activation causes the expression of proapoptotic genes with apoptosis showing both its upregulation and downregulation accordingly [64]. Increased levels of caspases, along with the hypoxic state, are involved in beta cell apoptosis [65]. Caspase-3, an important effector of the apoptotic pathways of DM, was also evaluated in our study [42]. A study among Caspase-3 knockout (Casp^−/−^) mice has shown that these mice were protected from the development of DM with a low-dose administration of STZ, which, otherwise, causes selective β cell destruction and further triggers the immune reactions in islet cells [42]. Studies with GLP-1 analogs among the animal models, in-vitro cell lines and human islet cells have shown a reduction in apoptosis, which was associated with a significant down-regulation of caspase-3 and up-regulation of bcl-2, and an increase in intracellular insulin content [47,64,66,67]. In an earlier study, Puerarin, one of the components of PTY-2, decreased the expression of caspase-3 in osteoblasts of diabetic rats and improved the pathological changes in diabetic rats [68].

HIF-1α is also a transcriptional activator of VEGF [69]. VEGF, a pro-angiogenic growth factor, helps in the vascularization of the pancreatic islets [31,70]. Currently, anti-VEGF therapy is approved for use in diabetic retinopathy [71]. Additionally, the effect of GLP-1 and DPP-IV inhibition are being evaluated in diabetic ulcers and for cardiovascular protective role [69,72]. Oxidative stress, measured by the presence of ROS, is also a promoter of angiogenesis [70].

Hyperglycemia also impairs nephrin signaling by increasing its internalization and the up-regulation of PKC-α expression, thus, playing an interesting role against gradual pancreatic β-cell loss in T2DM [29].

Currently, there is a need of antidiabetic agents with a wider spectrum of actions. As the roles of inflammation, oxidative stress, and apoptosis have become clearer over years, the currently available drugs should be re-evaluated for their effects on newer targets. Additionally, there is a need of newer agents which have action beyond the glucose-lowering potential. Various compounds have been studied for their role in the treatment of DM. A novel transcriptional inhibitor of TNF-α, MDL 201.449, has been found to reduce TNF-α mRNA levels dose dependently and prevent the development of hyperglycemia among mice following STZ injections [73].

PTY-2 is a herbal medicine under evaluation for its role in DM. As mentioned earlier, PTY-2 has shown anti-diabetic action by inhibition of DPP-IV enzyme, by acting as incretin receptor agonist, and by decreasing β cell apoptosis. Further pre-clinical and clinical research can help in the utilization of PTY-2 as a treatment option in DM. PTY-2 can be a less costly treatment option as compared to the already available anti-diabetic drugs. As PTY-2 extract is composed of many phytochemicals, it can be effective for multiple diseases. On the other hand, an evaluation of the limitations of the study shows that this study did not evaluate the role of individual phytochemicals in DM. Overall, further post-translational studies are required to completely understand the protective effect of PTY-2 on pancreas.

## Conclusion

Administration of PTY-2 for 10 days decreased the expressions of various biomarkers of oxidative stress and inflammation such as MMP-9, SOD, NF-κB, VEGF, TNF-α, Caspase-3, IL-6, and HIF-1α among STZ-induced diabetic rats as compared to controls. In addition, our results showed that PTY-2 may play a protective role in STZ-induced pancreatic stress. However, further research is needed to establish the role of PTY-2 in the treatment of DM.

## Abbreviations

STZ: Streptozotocon
PTY 2: *Pueraria tuberosa* water extract
SOD: Superoxide dismutase
NFkB: nuclear factor kappa-light-chain-enhancer of activated B cells
PKC ε: Protein kinase C ε
TNF α: Tumour necrosis factor α
MMP 9: Matrix metallopeptidase 9
HIF 1α: Hypoxia inducible factor α
VEGF: Vascular Endothelial Growth Factor
PTY 2: *Pueraria tuberosa* aqueous extract
MMP 9: Matrix metallopeptidase 9
SOD: Superoxide Dismutase
VEGF: Vascular Endothelial Growth Factor
TNF alpha: Tumor Necrosis Factor-alpha
IL 6: Interleukin 6
Hif-1α: Hypoxia-inducible factor 1-alpha
NF-κB: Nuclear Factor kappa-light-chain-enhancer of activated B cells
DM: Diabetes Mellitus
DPP-IV: Dipeptidyl Peptidase 4
GLP-1: Glucagon-like peptide 1
GIP: Glucose-Dependent Insulinotropic Peptide

## Declarations

### Ethics approval and consent to participate

The protocol was approved by the Institute Ethical Committee (Dean/2015/CAEC/1266), Institute of Medical Sciences, Banaras Hindu University.

### Consent for publication

Not Applicable

### Availability of data and materials

The data and materials supporting the conclusions of this work are included in the article.

## Authors’ contributions

Shivani Srivastava conceived, designed, implemented and analyzed the data for all the experiments and wrote the manuscript. Tissue slides were prepared by Harsh Pandey. Yamini Bhusan Tripathi and Surya Kumar Singh provided guidance for all levels from conception to manuscript writing.

## Acknowledgments

We are thankful to the Head of the Department, Dr. Amrita G. Kar, and all laboratory technicians of the Department of Pathology, IMS, BHU for their help in IHC slide preparations. Heartfelt thanks to Miss. Surabhi Singh, Dr. Mukulika Ray, Dr. Deoprakash Chaturvedi & Prof. Subhash Lakhotia, Department of Zoology, BHU for their supportive help in Confocal Microscopy. We are highly thankful to ISLS, BHU for providing us the facility to perform Confocal Microscopy. Last but not least, we are grateful to Mrs. Durgavati Yadav, Mrs. Rashmi Shukla and Prerna Aditi for helping us in sample processing.

## References

[1] WHO. WHO | India: first to adapt the Global Monitoring Framework on noncommunicable diseases (NCDs). WHO 2015.

[2] Yadav D, Lowenfels AB. The epidemiology of pancreatitis and pancreatic cancer. Gastroenterology 2013;144:1252–61. doi:10.1053/j.gastro.2013.01.068.

[3] DiMagno MJ, DiMagno EP. Chronic pancreatitis. Curr Opin Gastroenterol 2012;28:523–31. doi:10.1097/MOG.0b013e3283567dea.

[4] Alsamarrai A, Das SLM, Windsor JA, Petrov MS. Factors That Affect Risk for Pancreatic Disease in the General Population: A Systematic Review and Meta-analysis of Prospective Cohort Studies. Clin Gastroenterol Hepatol 2014;12:1635–1644.e5. doi:10.1016/j.cgh.2014.01.038.

[5] Das SLM, Singh PP, Phillips ARJ, Murphy R, Windsor JA, Petrov MS. Newly diagnosed diabetes mellitus after acute pancreatitis: a systematic review and meta-analysis. Gut 2014;63:818–31. doi:10.1136/gutjnl-2013-305062.

[6] Gupta S, Vittinghoff E, Bertenthal D, Corley D, Shen H, Walter LC, et al. New-Onset Diabetes and Pancreatic Cancer. Clin Gastroenterol Hepatol 2006;4:1366–72. doi:10.1016/j.cgh.2006.06.024.

[7] Ben Q, Xu M, Ning X, Liu J, Hong S, Huang W, et al. Diabetes mellitus and risk of pancreatic cancer: A meta-analysis of cohort studies. Eur J Cancer 2011;47:1928–37. doi:10.1016/j.ejca.2011.03.003.

[8] Perrin MC, Terry MB, Kleinhaus K, Deutsch L, Yanetz R, Tiram E, et al. Gestational diabetes as a risk factor for pancreatic cancer: a prospective cohort study. BMC Med 2007;5:25. doi:10.1186/1741-7015-5-25.

[9] Girman CJ, Kou TD, Cai B, Alexander CM, O’Neill EA, Williams-Herman DE, et al. Patients with type 2 diabetes mellitus have higher risk for acute pancreatitis compared with those without diabetes. Diabetes, Obes Metab 2010;12:766–71. doi:10.1111/j.1463-1326.2010.01231.x.

[10] Rosendahl J, Bödeker H, Mössner J, Teich N. Hereditary chronic pancreatitis. Orphanet J Rare Dis 2007;2:1. doi:10.1186/1750-1172-2-1.

[11] Srivastava S, Shree P, Pandey H, Tripathi YB. Incretin hormones receptor signaling plays the key role in antidiabetic potential of PTY-2 against STZ-induced pancreatitis. Biomed Pharmacother 2018;97:330–8. doi:10.1016/j.biopha.2017.10.071.

[12] Srivastava S, Koley TK, Singh SK, Tripathi YB. THE TUBER EXTRACT OF PUERARIA TUBEROSA LINN. COMPETITIVELY INHIBITS DPP-IV ACTIVITY IN NORMOGLYCEMIC RATS. Int J Pharm Pharm Sci 2015;7:7–11.

[13] Srivastava S, Shree P, Tripathi YB. Active phytochemicals of Pueraria tuberosa for DPP-IV inhibition: in silico and experimental approach. J Diabetes Metab Disord 2017;16:46. doi:10.1186/s40200-017-0328-0.

[14] Tripathi AK, Kohli S. Anti-Diabetic Activity and Phytochemical Screening of Crude Extracts of PuerariaTuberosa DC. (FABACEAE) Grown in India on STZ-Induced Diabetic Rats. Asian J Med Pharm Res 2013;3:66–73.

[15] Pandey N, Yadav D, Pandey V, Tripathi YB. Anti-inflammatory effect of Pueraria tuberosa extracts through improvement in activity of red blood cell anti-oxidant enzymes. Ayu 2013;34:297–301. doi:10.4103/0974-8520.123131.

[16] Pandey N, Chaurasia JK, Tiwari OP, Tripathi YB. Antioxidant properties of different fractions of tubers from Pueraria tuberosa Linn. Food Chem 2007;105:219–22. doi:10.1016/j.foodchem.2007.03.072.

[17] Tripathi YB, Nagwani S, Mishra P, Jha A, Rai SP. Protective effect of Pueraria tuberosa DC. embedded biscuit on cisplatin-induced nephrotoxicity in mice. J Nat Med 2012;66:109–18. doi:10.1007/s11418-011-0559-1.

[18] Yadav D, Kumar M, Tripathi YB. Methanolic extract of tubers of Pueraria tuberosa Linn. ameliorates glycerol induced acute kidney injury in rats. J Chem Pharm Res 2016;8:133–9.

[19] Verma SK, Jain V, Singh DP. Effect of Pueraria tuberosa DC. (Indian Kudzu) on Blood Pressure, Fibrinolysis and Oxidative Stress in Patients with Stage 1 Hypertension. Pakistan J Biol Sci 2012;15:742–7. doi:10.3923/pjbs.2012.742.747.

[20] Pramanik SS, Sur TK, Debnath PK, Bhattacharyya D. Effect of Pueraria tuberosa tuber extract on chronic foot shock stress in Wistar rats. Nepal Med Coll J 2010;12:234–8.

[21] Burke SJ, Lu D, Sparer TE, Karlstad MD, Collier JJ. Transcription of the gene encoding TNF-α is increased by IL-1β in rat and human islets and β-cell lines. Mol Immunol 2014;62:54–62. doi:10.1016/j.molimm.2014.05.019.

[22] Uemura S, Matsushita H, Li W, Glassford AJ, Asagami T, Lee KH, et al. Diabetes mellitus enhances vascular matrix metalloproteinase activity: role of oxidative stress. Circ Res 2001;88:1291–8.

[23] Shibaji T, Nagao M, Ikeda N, Kanehiro H, Hisanaga M, Ko S, et al. Prognostic significance of HIF-1 alpha overexpression in human pancreatic cancer. Anticancer Res n.d.;23:4721–7.

[24] Pini M, Rhodes DH, Castellanos KJ, Hall AR, Cabay RJ, Chennuri R, et al. Role of IL-6 in the resolution of pancreatitis in obese mice. J Leukoc Biol 2012;91:957–66. doi:10.1189/jlb.1211627.

[25] Reinert RB, Brissova M, Shostak A, Pan FC, Poffenberger G, Cai Q, et al. Vascular Endothelial Growth Factor-A and Islet Vascularization Are Necessary in Developing, but Not Adult, Pancreatic Islets. Diabetes 2013;62:4154–64. doi:10.2337/db13-0071.

[26] Singh N, Bhardwaj P, Pandey RM, Saraya A. Oxidative stress and antioxidant capacity in patients with chronic pancreatitis with and without diabetes mellitus. Indian J Gastroenterol 2012;31:226–31. doi:10.1007/s12664-012-0236-7.

[27] Dabhi B, Mistry KN. Oxidative stress and its association with TNF-α-308 G/C and IL-1α-889 C/T gene polymorphisms in patients with diabetes and diabetic nephropathy. Gene 2015;562:197–202. doi:10.1016/j.gene.2015.02.069.

[28] Putaala H, Soininen R, Kilpeläinen P, Wartiovaara J, Tryggvason K. The murine nephrin gene is specifically expressed in kidney, brain and pancreas: inactivation of the gene leads to massive proteinuria and neonatal death. Hum Mol Genet 2001;10:1–8.

[29] Kapodistria K, Tsilibary E-P, Politis P, Moustardas P, Charonis A, Kitsiou P. Nephrin, a transmembrane protein, is involved in pancreatic beta-cell survival signaling. Mol Cell Endocrinol 2015;400:112–28. doi:10.1016/j.mce.2014.11.003.

[30] Watada H. Role of VEGF-A in pancreatic beta cells. Endocr J 2010;57:185–91.

[31] Brissova M, Shostak A, Shiota M, Wiebe PO, Poffenberger G, Kantz J, et al. Pancreatic Islet Production of Vascular Endothelial Growth Factor-A Is Essential for Islet Vascularization, Revascularization, and Function. Diabetes 2006;55:2974–85. doi:10.2337/db06-0690.

[32] Reinert RB, Cai Q, Hong J-Y, Plank JL, Aamodt K, Prasad N, et al. Vascular endothelial growth factor coordinates islet innervation via vascular scaffolding. Development 2014;141:1480–91. doi:10.1242/dev.098657.

[33] Zhen G-D, Zhao L-B, Wu S-S, Chen M-Y, Li Z-H, Zhou S-Z, et al. Associations of MMP-2 and MMP-9 gene polymorphism with ulinastatin efficacy in patients with severe acute pancreatitis. Biosci Rep 2017;37:BSR20160612. doi:10.1042/BSR20160612.

[34] De Palma AM, Verbeken E, Aelst I Van, Van den Steen PE, Opdenakker G, Neyts J. Increased gelatinase B/matrix metalloproteinase 9 (MMP-9) activity in a murine model of acute coxsackievirus B4-induced pancreatitis. Virology 2008;382:20–7. doi:10.1016/j.virol.2008.08.046.

[35] Opdenakker G, Van den Steen PE, Van Damme J. Gelatinase B: a tuner and amplifier of immune functions. Trends Immunol 2001;22:571–9.

[36] Rakonczay Z, Hegyi P, Takacs T, McCarroll J, Saluja AK. The role of NF-B activation in the pathogenesis of acute pancreatitis. Gut 2008;57:259–67. doi:10.1136/gut.2007.124115.

[37] Patel S, Santani D. Role of NF-kappa B in the pathogenesis of diabetes and its associated complications. Pharmacol Rep n.d.;61:595–603.

[38] Swaroop JJ, Rajarajeswari D, Naidu JN. Association of TNF-α with insulin resistance in type 2 diabetes mellitus. Indian J Med Res 2012;135:127–30.

[39] Cantley J, Burchfield JG, Pearson GL, Schmitz-Peiffer C, Leitges M, Biden TJ. Deletion of PKC◻ Selectively Enhances the Amplifying Pathways of Glucose-Stimulated Insulin Secretion via Increased Lipolysis in Mouse ɴ_ʟ_-Cells n.d. doi:10.2337/db09-0132.

[40] Kim M-J, Lee Y-S, Lee K-H, Min DS, Yoon S-H, Hahn SJ, et al. Site-Specific Localization of Protein Kinase C Isoforms in Rat Pancreas. Pancreatology 2001;1:36–42. doi:10.1159/000055790.

[41] Moritz W, Meier F, Stroka DM, Giuliani M, Kugelmeier P, Nett PC, et al. Apoptosis in hypoxic human pancreatic islets correlates with HIF-1α expression. FASEB J 2002;16:745–7. doi:10.1096/fj.01-0403fje.

[42] Liadis N, Murakami K, Eweida M, Elford AR, Sheu L, Gaisano HY, et al. Caspase-3-Dependent -Cell Apoptosis in the Initiation of Autoimmune Diabetes Mellitus. Mol Cell Biol 2005;25:3620–9. doi:10.1128/MCB.25.9.3620-3629.2005.

[43] Asthana S, Agarwal T, Singothu S, Samal A, Banerjee I, Pal K, et al. Molecular Docking and Interactions of Pueraria Tuberosa with Vascular Endothelial Growth Factor Receptors. Indian J Pharm Sci 2015;77:439–45.

[44] Pandey N, Tripathi YB. Antioxidant activity of tuberosin isolated from Pueraria tuberose Linn. J Inflamm (Lond) 2010;7:47. doi:10.1186/1476-9255-7-47.

[45] Cieŝlak M, Wojtczak A, Cieŝlak M. Role of pro-inflammatory cytokines of pancreatic islets and prospects of elaboration of new methods for the diabetes treatment n.d. doi:10.18388/abp.2014_853.

[46] Lee Y-S, Jun H-S. Anti-Inflammatory Effects of GLP-1-Based Therapies beyond Glucose Control. Mediators Inflamm 2016;2016:1–11. doi:10.1155/2016/3094642.

[47] Farilla L, Hui H, Bertolotto C, Kang E, Bulotta A, Di Mario U, et al. Glucagon-Like Peptide-1 Promotes Islet Cell Growth and Inhibits Apoptosis in Zucker Diabetic Rats. Endocrinology 2002;143:4397–408. doi:10.1210/en.2002-220405.

[48] Descamps FJ, Martens E, Ballaux F, Geboes K, Opdenakker G. In vivo activation of gelatinase B/MMP-9 by trypsin in acute pancreatitis is a permissive factor in streptozotocin-induced diabetes. J Pathol 2004;204:555–61. doi:10.1002/path.1669.

[49] Descamps FJ, Van den Steen PE, Martens E, Ballaux F, Geboes K, Opdenakker G. Gelatinase B is diabetogenic in acute and chronic pancreatitis by cleaving insulin. FASEB J 2003;17:887–9. doi:10.1096/fj.02-0725fje.

[50] Ceriello A, La Sala L, De Nigris V, Pujadas G, Rondinelli M, Genovese S. GLP-1 reduces metalloproteinase-9 induced by both hyperglycemia and hypoglycemia in type 1 diabetes. The possible role of oxidative stress. Ther Clin Risk Manag 2015;11:901–3. doi:10.2147/TCRM.S83322.

[51] Christoffersson G, Waldén T, Sandberg M, Opdenakker G, Carlsson P-O, Phillipson M. Matrix Metalloproteinase-9 Is Essential for Physiological Beta Cell Function and Islet Vascularization in Adult Mice. Am J Pathol 2015;185:1094–103. doi:10.1016/j.ajpath.2014.12.009.

[52] Mack E, Ziv E, Reuveni H, Kalman R, Niv MY, Jörns A, et al. Prevention of insulin resistance and beta-cell loss by abrogating PKCε-induced serine phosphorylation of muscle IRS-1 in *Psammomys obesus*. Diabetes Metab Res Rev 2008;24:577–84. doi:10.1002/dmrr.881.

[53] Croft M, Benedict CA, Ware CF. Clinical targeting of the TNF and TNFR superfamilies. Nat Rev Drug Discov 2013;12:147–68. doi:10.1038/nrd3930.

[54] Chen W, Zheng Z, Duan J, Wang X, Wu S, Wang W, et al. Quantitation of nuclear factor kappa B activation in pancreatic acinar cells during rat acute pancreatitis by flow cytometry. Int J Clin Exp Med 2015;8:10143–51.

[55] Steinle AU, Weidenbach H, Wagner M, Adler G, Schmid RM. NF-κB/Rel activation in cerulein pancreatitis. Gastroenterology 1999;116:420–30. doi:10.1016/S0016-5085(99)70140-X.

[56] Algül H, Treiber M, Lesina M, Nakhai H, Saur D, Geisler F, et al. Pancreas-specific RelA/p65 truncation increases susceptibility of acini to inflammation-associated cell death following cerulein pancreatitis. J Clin Invest 2007;117:1490–501. doi:10.1172/JCI29882.

[57] Fantini L, Tomassetti P, Pezzilli R. Management of acute pancreatitis: current knowledge and future perspectives. World J Emerg Surg 2006;1:16. doi:10.1186/1749-7922-1-16.

[58] Hu X, Liu S, Liu X, Zhang J, Liang Y, Li Y. DPP-4 (CD26) inhibitor sitagliptin exerts anti-inflammatory effects on rat insulinoma 26 (RINm) cells via suppressing NF-κB activation. Endocrine 2017;55:754–63. doi:10.1007/s12020-016-1073-8.

[59] Liuwantara D, Elliot M, Smith MW, Yam AO, Walters SN, Marino E, et al. Nuclear factor-kappaB regulates beta-cell death: a critical role for A20 in beta-cell protection. Diabetes 2006;55:2491–501. doi:10.2337/db06-0142.

[60] Mandrup-Poulsen T. Apoptotic signal transduction pathways in diabetes. Biochem Pharmacol 2003;66:1433–40.

[61] De M. Potential role of TNF-alpha in the pathogenesis of insulin resistance and type 2 diabetes. Trends Endocrinol Metab 2000;11:21–7.

[62] Catrina S-B, Okamoto K, Pereira T, Brismar K, Poellinger L. Hyperglycemia regulates hypoxia-inducible factor-1alpha protein stability and function. Diabetes 2004;53:3226–32.

[63] Bento CF, Pereira P. Regulation of hypoxia-inducible factor 1 and the loss of the cellular response to hypoxia in diabetes. Diabetologia 2011;54:1946–56. doi:10.1007/s00125-011-2191-8.

[64] Velmurugan K, Balamurugan AN, Loganathan G, Ahmad A, Hering BJ, Pugazhenthi S. Antiapoptotic Actions of Exendin-4 against Hypoxia and Cytokines Are Augmented by CREB. Endocrinology 2012;153:1116–28. doi:10.1210/en.2011-1895.

[65] Greijer AE, van der Wall E. The role of hypoxia inducible factor 1 (HIF-1) in hypoxia induced apoptosis. J Clin Pathol 2004;57:1009–14. doi:10.1136/jcp.2003.015032.

[66] Farilla L, Bulotta A, Hirshberg B, Li Calzi S, Khoury N, Noushmehr H, et al. Glucagon-Like Peptide 1 Inhibits Cell Apoptosis and Improves Glucose Responsiveness of Freshly Isolated Human Islets. Endocrinology 2003;144:5149–58. doi:10.1210/en.2003-0323.

[67] Li Y, Hansotia T, Yusta B, Ris F, Halban PA, Drucker DJ. Glucagon-like Peptide-1 Receptor Signaling Modulates β Cell Apoptosis. J Biol Chem 2003;278:471–8. doi:10.1074/jbc.M209423200.

[68] Liang J, Chen H, Pan W, Xu C. Puerarin inhibits caspase-3 expression in osteoblasts of diabetic rats. Mol Med Rep 2012;5:1419–22. doi:10.3892/mmr.2012.854.

[69] Marfella R, Sasso FC, Rizzo MR, Paolisso P, Barbieri M, Padovano V, et al. Dipeptidyl peptidase 4 inhibition may facilitate healing of chronic foot ulcers in patients with type 2 diabetes. Exp Diabetes Res 2012;2012:892706. doi:10.1155/2012/892706.

[70] El-Refaei MF, Abduljawad SH, Alghamdi AH. Alternative Medicine in Diabetes - Role of Angiogenesis, Oxidative Stress, and Chronic Inflammation. Rev Diabet Stud 2014;11:231–44. doi:10.1900/RDS.2014.11.231.

[71] Osaadon P, Fagan XJ, Lifshitz T, Levy J. A review of anti-VEGF agents for proliferative diabetic retinopathy. Eye 2014;28:510–20. doi:10.1038/eye.2014.13.

[72] Xiao-Yun X, Zhao-Hui M, Ke C, Hong-Hui H, Yan-Hong X. Glucagon-like peptide-1 improves proliferation and differentiation of endothelial progenitor cells via upregulating VEGF generation. Med Sci Monit 2011;17:BR35–41.

[73] Holstad M, Sandler S. A Transcriptional Inhibitor of TNF-α Prevents Diabetes Induced by Multiple Low-Dose Streptozotocin Injections in Mice. J Autoimmun 2001;16:441–7. doi:10.1006/jaut.2001.0506.

